# Deconstructing isolation-by-distance: the genomic consequences of limited dispersal

**DOI:** 10.1101/093989

**Authors:** Stepfanie M. Aguillon, John W. Fitzpatrick, Reed Bowman, Stephan J. Schoech, Andrew G. Clark, Graham Coop, Nancy Chen

## Abstract

Geographically limited dispersal can shape genetic population structure and result in a correlation between genetic and geographic distance, commonly called isolation-by-distance. Despite the prevalence of isolation-by-distance in nature, to date few studies have empirically demonstrated the processes that generate this pattern, largely because few populations have direct measures of individual dispersal and pedigree information. Intensive, long-term demographic studies and exhaustive genomic surveys in the Florida Scrub-Jay (*Aphelocoma coerulescens*) provide an excellent opportunity to investigate the influence of dispersal on genetic structure. Here, we used a panel of genome-wide SNPs and extensive pedigree information to explore the role of limited dispersal in shaping patterns of isolation-by-distance in both sexes, and at an exceedingly fine spatial scale (within ~10 km). Isolation-by-distance patterns were stronger in male-male and male-female comparisons than in female-female comparisons, consistent with observed differences in dispersal propensity between the sexes. Using the pedigree, we demonstrated how various genealogical relationships contribute to fine-scale isolation-by-distance. Simulations using field-observed distributions of male and female natal dispersal distances showed good agreement with the distribution of geographic distances between breeding individuals of different pedigree relationship classes. Furthermore, we extended Malécot’s theory of isolation-by-distance by building coalescent simulations parameterized by the observed dispersal curve, population density, and immigration rate, and showed how incorporating these extensions allows us to accurately reconstruct observed sex-specific isolation-by-distance patterns in autosomal and Z-linked SNPs. Therefore, patterns of fine-scale isolation-by-distance in the Florida Scrub-Jay can be well understood as a result of limited dispersal over contemporary timescales.

**Author Summary:** Dispersal is a fundamental component of the life history of most organisms and therefore influences many biological processes. Dispersal is particularly important in creating genetic structure on the landscape. We often observe a pattern of decreased genetic relatedness between individuals as geographic distances increases, or isolation-by-distance. This pattern is particularly pronounced in organisms with extremely short dispersal distances. Despite the ubiquity of isolation-by-distance patterns in nature, there are few examples that explicitly demonstrate how limited dispersal influences spatial genetic structure. Here we investigate the processes that result in spatial genetic structure using the Florida Scrub-Jay, a bird with extremely limited dispersal behavior and extensive genome-wide data. We take advantage of the long-term monitoring of a contiguous population of Florida Scrub-Jays, which has resulted in a detailed pedigree and measurements of dispersal for hundreds of individuals. We show how limited dispersal results in close genealogical relatives living closer together geographically, which generates a strong pattern of isolation-by-distance at an extremely small spatial scale (<10 km) in just a few generations. Given the detailed dispersal, pedigree, and genomic data, we can achieve a fairly complete understanding of how dispersal shapes patterns of genetic diversity over short spatial scales.

## Introduction

The movement of individuals over the landscape (dispersal) influences biological processes and diversity at many levels [1], ranging from interactions between individuals to persistence of populations or species over time [2-4]. Limited dispersal is also central to generating and maintaining spatial genetic structure within species. In particular, geographically-limited dispersal can result in isolation-by-distance, a pattern of increased genetic differentiation [5, 6] or, conversely, decreased genetic relatedness [7-9] between individuals as geographic distance increases. This pattern results because genetic drift can act to differentiate allele frequencies faster than dispersal can homogenize them among geographically distant populations. A correlation between genetic differentiation and geographic distance is observed in many empirical systems, consistent with isolation-by-distance being an important process in structuring genetic diversity [10, 11].

Despite the fact that correlations between genetic differentiation and geographic distance are common across many types of organisms, to date, there are few existing empirical demonstrations of how contemporary patterns of dispersal generate spatial patterns of genetic variation and contribute to observed patterns of isolation-by-distance. This is, in part, because dispersal is hard to estimate empirically, as it requires monitoring many individuals over long periods of time across the full range of potential dispersal distances [12]. In addition, the effective population density of reproducing individuals must be known in order to parameterize genetic drift in models of isolation-by-distance. Therefore, in practice it is hard to know whether the observation of isolation-by-distance is truly consistent with contemporary patterns of dispersal. Indeed, many studies use genetic isolation-by-distance patterns to infer dispersal distances, as a substitute for the more difficult exercise of measuring dispersal directly in the field [13-17]. A second issue is that in most studied systems, populations are compared over large spatial scales, so the pattern of isolation-by-distance reflects the dynamics of genetic drift and dispersal over tens of thousands of generations. These empirical patterns reflect large-scale population movements (*e.g.*, expansions from glacial refugia [18]) that may not reflect the equilibrium outcome of individual dispersal and genetic drift. While studies have reported fine-scale population structure [19-25], it has been difficult to deconstruct these patterns to understand what mechanisms actually create them.

Patterns of isolation-by-distance can reflect underlying biological processes. Since the early development of the isolation-by-distance theory, differences in mating systems and dispersal propensity have both been known to generate differences in isolation-by-distance patterns [5]. In many organisms, dispersal often differs between the sexes: males tend to disperse farther in mammals (male-biased dispersal), but females tend to disperse farther in birds (female-biased dispersal) [26, 27]. When dispersal patterns differ between the sexes, the less dispersive sex tends to have stronger overall genetic structure than the more dispersive sex [21, 28, 29]. Similarly, sex-biased dispersal is expected to result in different levels of genetic structure in markers with different inheritance patterns. For example, in a system where females are both the heterogametic sex (*e.g.*, in birds, females are ZW and males are ZZ) and more dispersive, autosomes may exhibit higher genetic differentiation than maternally inherited markers (*e.g.*, mitochondrial DNA), but lower genetic differentiation than the Z chromosome [16, 30, 31].

Here we examine the causes of fine-scale isolation-by-distance in a non-migratory bird, the Florida Scrub-Jay (*Aphelocoma coerulescens*), based on a long-term population study that has yielded high-quality genetic and pedigree information for many individuals, as well as particularly detailed information on individual dispersal distances. Florida Scrub-Jays have limited, female-biased natal dispersal, and individuals essentially never move once established as a breeding adult [32, 33]. A population of Florida Scrub-Jays at Archbold Biological Station in central Florida has been the focus of intense monitoring since 1969, resulting in observed natal dispersal distances for hundreds of individuals and an extensive pedigree [32, 34, 35]. Moreover, nearly all nestlings and breeders present in the population during the past two decades were genotyped in a recent study [36]. These long-term dispersal, pedigree, and genomic data make the Florida Scrub-Jay an unusually tractable study system in which to explore how dispersal influences patterns of isolation-by-distance.

Previous work on Florida Scrub-Jays using microsatellite markers has shown isolation-by-distance across multiple populations [3]. Here, we present evidence for fine-scale isolation-by-distance within a single contiguous population of Florida Scrub-Jays, and combine genetic, pedigree, and dispersal information to reveal how patterns of isolation-by-distance are created in nature. We find more isolation-by-distance in males than in females, corresponding to predicted differences resulting from female-biased dispersal patterns. We break down our data into pedigree relationships to demonstrate that isolation-by-distance is a consequence of close relatives living geographically close together. We perform simulations that successfully reconstruct the empirical distances between individuals of different kinship classes using only the dispersal curves. Finally, we use extensive coalescent simulations parameterized by the dispersal curve, population density, and immigration rate to yield an excellent fit to observed isolation-by-distance patterns.

## Results/Discussion

### Limited dispersal results in isolation-by-distance at small spatial scales

We documented natal dispersal distances for 382 male and 290 female Florida Scrub-Jays that were born and established as breeders within the population at Archbold Biological Station between 1990-2013. Dispersal curves for both males and females were strongly leptokurtic, consistent with previous studies (Fig 1A; [3, 34]). Here we considered only dispersal within the Archbold population; therefore, our dispersal curves do not capture any long-distance dispersal events, which occur rarely [3]. Females disperse significantly farther than males, with a median ± SE distance of 1,149 ± 108 m and 488 ± 43 m, respectively (Wilcoxon rank sum test, *p* < 2.2 x 10^−16^). Florida Scrub-Jays disperse extremely short distances compared with other bird species [34, 37]. The shorter male dispersal distances compared with females may be due in part to differences in territory acquisition between the sexes. Florida Scrub-Jay males are able to acquire breeding territories through budding from the parental territory or inheritance of the parental territory [32], while territory budding and inheritance is extremely rare in females [34].

**Fig 1.**
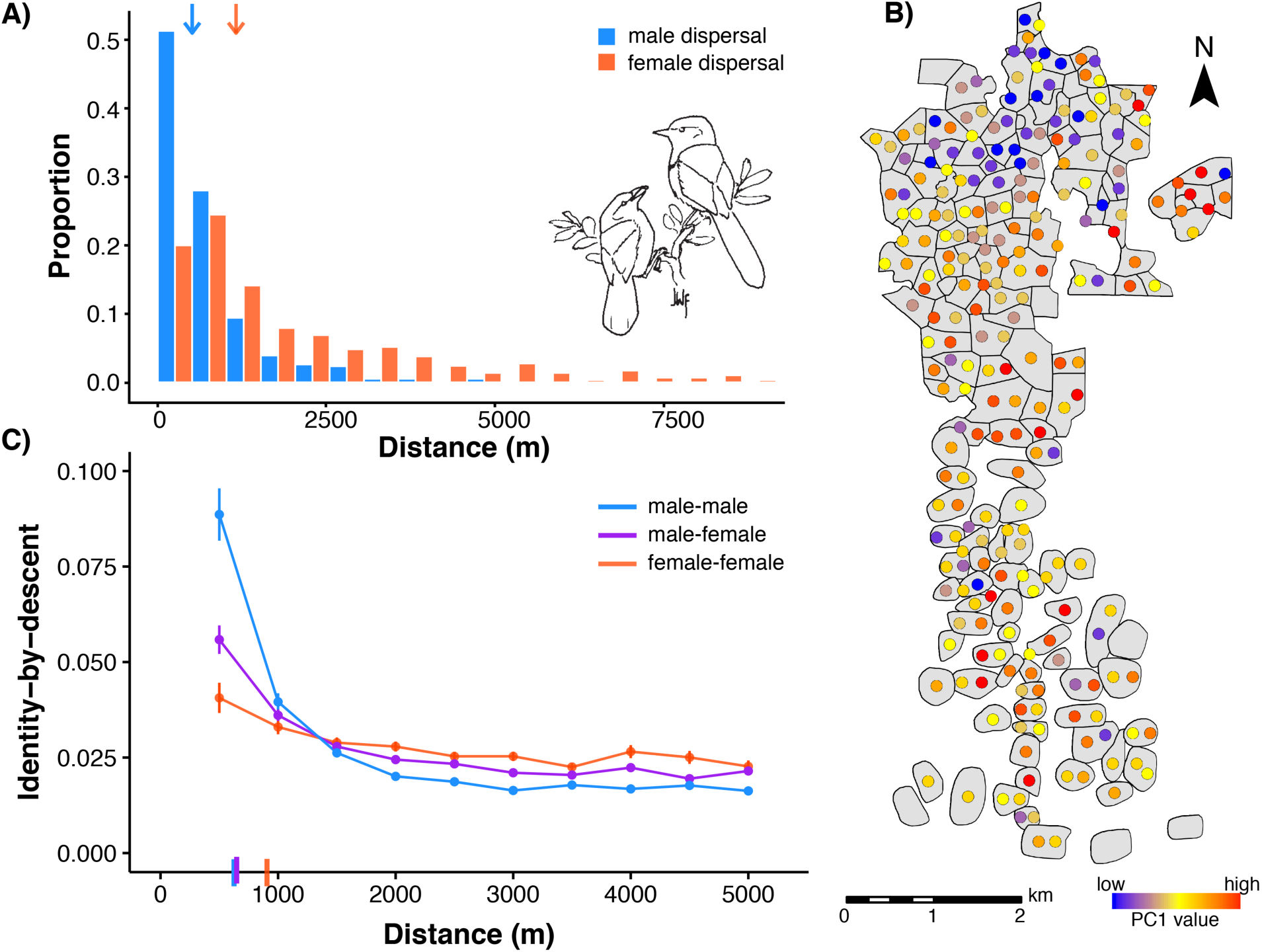
Dispersal curves and isolation-by-distance patterns in the Florida Scrub-Jay. (A) Natal dispersal distances for Florida Scrub-Jays born and breeding within Archbold Biological Station (1990-2013, *n* = 672) are significantly shorter in males (blue bars; median ± SE = 488 ± 43 m) than in females (salmon bars; 1,149 ± 108 m; Wilcoxon rank sum test, *p* < 2.2 x 10^−16^). Median values are shown with arrows at top of plot. Florida Scrub-Jay drawing by JWF. (B) Map of breeding territories (gray polygons) for a representative year (2008) within Archbold with individual breeders colored by PC1 values shows isolation-by-distance from north to south. (C) Isolation-by-distance patterns in autosomal SNPs shown with standard error bars. The decline in identity-by-descent with geographic distance is stronger in male-male (blue) and male-female (purple) pairwise comparisons than in female-female comparisons (salmon). δ values, the distance where identity-by-descent drops halfway to the mean (see text for details), are shown as dashes on the x-axis.

To explore the genetic implications of this limited, sex-biased dispersal, we genotyped all breeding adults in the Archbold population in 2003, 2008, and 2013 (*n* = 513) at 7,843 autosomal SNPs and 277 Z-linked SNPs [36]. We conducted principal component analysis (PCA) separately for all breeding adults, male breeders, and female breeders to visually summarize patterns of autosomal genetic variation within the population. We see genetic differentiation along the north/south axis of Archbold in the first two PC axes when we map breeders to their breeding territories (Fig 1B, S1 Fig). Indeed, the top two principal components (PC1 and PC2, 14.6% and 13.1% of the variation, respectively) are significantly correlated with north-south position under the Universal Transverse Mercator coordinate system (henceforth “UTM northing”; S1 Table). We found significant correlations with UTM northing for both PC1 and PC2 in males, but only PC1 is significantly correlated with UTM northing in females (S2 Fig, S1 Table). Correlation coefficients for PC1 with UTM northing are higher in males than in females (S1 Table). This fine-scale spatial structure is likely a direct result of the unusually limited natal dispersal and female-biased dispersal of these birds (Fig 1A; overall median ± SE = 647 ± 57 m).

To test for isolation-by-distance, we quantified autosomal genetic relatedness between all possible pairs of individuals in the dataset as the estimated proportion of the genome shared identical-by-descent. Under a model of isolation-by-distance, the proportion of the genome shared identical-by-descent should decrease as the distance between individuals in a pair increases. Plotting genetic relatedness against geographic distance for all unique pairs across all years, we found a clear pattern of isolation-by-distance (Fig 1C) at a fine spatial scale (Archbold is ~10 km from north to south; Fig 1B). We used Mantel correlograms to compare pairwise geographic and genetic distances (identity-by-descent) within distinct distance class bins across all pairwise comparisons, all male-male pairs, and all female-female pairs. Mantel correlograms are useful for testing spatial genetic structure when the relationship between geographic and genetic distance is exponential-like rather than linear [38, 39]. We found significant correlations at more distance classes in all breeders and male-male pairs than in female-female pairs (S2 Table), which indicates stronger patterns of isolation-by-distance in males than in females, consistent with the observed female-biased dispersal.

To measure the strength of isolation-by-distance in different subsets of the data, we fitted loess curves and used them to estimate the distance (δ) where the proportion of the genome shared identical-by-descent drops halfway to the mean from its maximum value. A lower δ indicates a more rapid decay of genetic relatedness by geographic distance, *i.e.,* more isolation-by-distance. We bootstrapped pairs of individuals to obtain 95% confidence intervals (CI) to assess significance and found stronger isolation-by-distance patterns in male-male (δ = 620 m, 95% CI = [604, 631]) and male-female comparisons (δ = 645 m, [622, 665]) than in female-female comparisons (δ = 903 m, [741, 1261]; Fig 1C), which is consistent with the strongly female-biased dispersal observed in this system.

Because of the detailed pedigree information available for the Florida Scrub-Jay population within Archbold, we have a rare opportunity to decompose the isolation-by-distance patterns found in this population by familial relationship. The Florida Scrub-Jay pedigree from our study population consists of 12,738 unique individuals over 14 generations and is largely complete (see S3 Table for a summary of the pedigree); here we identify relationships up to fourth cousins. For each pair of individuals in our dataset, we extracted their closest genealogical relationship from the pedigree (*e.g.*, 1,532 of 130,618 pairs have a relationship closer than first cousins; S3 Table) and calculated the pedigree-based coefficient of relationship (*r*). We plotted identity-by-descent for pairs of individuals against the geographic distance between those individuals, coloring points by their pedigree relationship (Fig 2). These plots clearly illustrate how isolation-by-distance results, in part, from closely related individuals, such as parent-offspring and full-siblings, remaining physically close together as breeders within neighborhoods of contiguous territories (Fig 2). The stronger signal of isolation-by-distance in male-male comparisons (Fig 2A) seems to be driven by the particularly short geographic distances between individuals in the highest pedigree relatedness classes (*e.g.*, parent-offspring, full-siblings, grandparent-grandchild, half-siblings, and aunt/uncle-nibling [“nibling” is a gender-neutral term for niece and nephew]).

**Fig 2.**
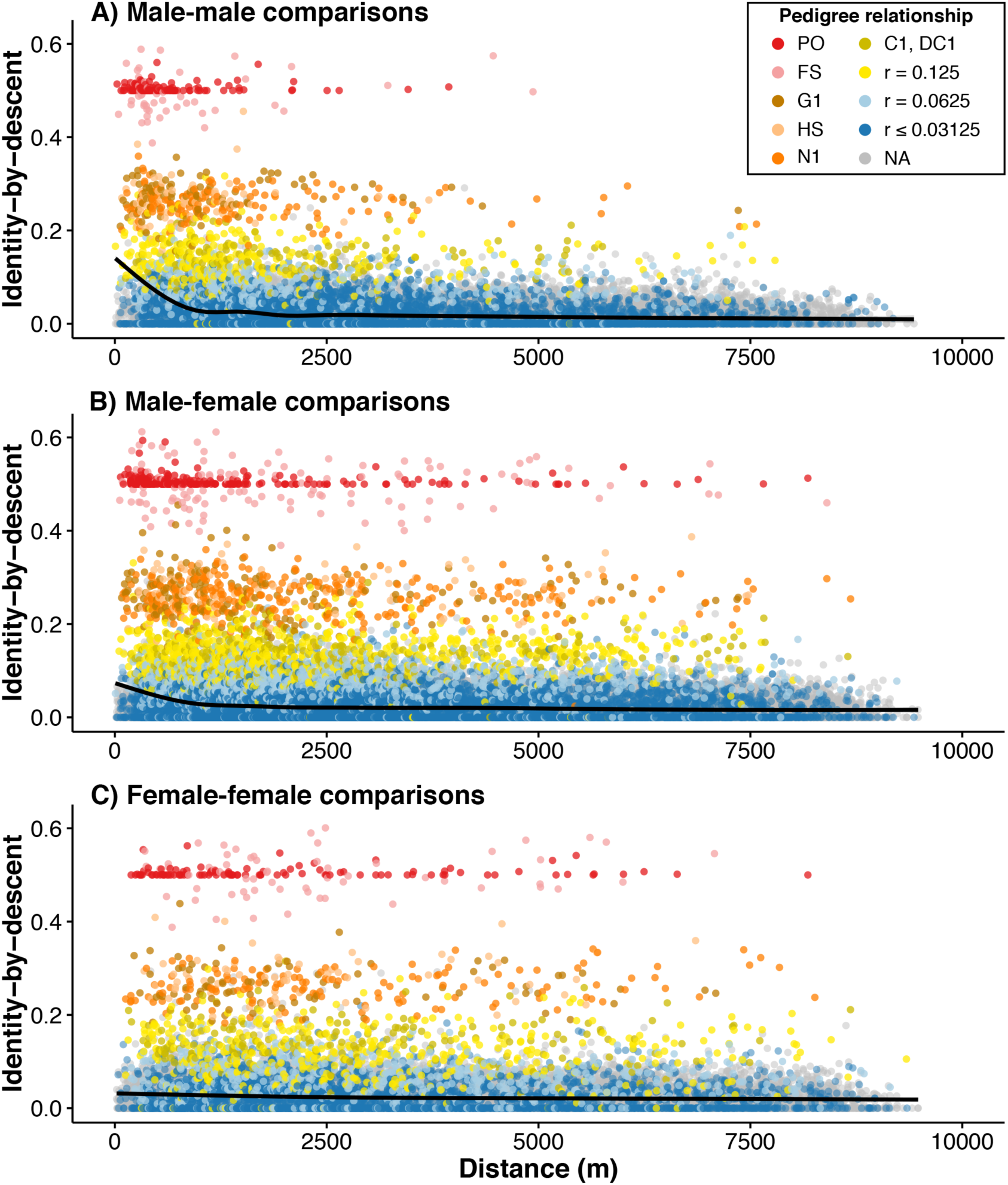
Isolation-by-distance patterns in Florida Scrub-Jays can be deconstructed by pedigree relatedness. Distance versus identity-by-descent in autosomal SNPs for all possible (A) male-male, (B) male-female, and (C) female-female comparisons is, in part, generated by highly related individuals remaining physically close together. Loess curves are shown in each panel. Isolation-by-distance patterns are significantly stronger in male-male (A) and male-female (B) comparisons than in female-female (C) comparisons. Points are colored by specific pedigree relationship or, for more distant relationships, grouped into a single coefficient of relationship (r) class. Gray points indicate no known pedigree relationship. Pedigree relationship abbreviations: PO = parent-offspring, FS = full-siblings, G1 = grandparent-grandchild, HS = half-siblings, N1 = aunt/uncle-nibling, C1 = first cousins, DC1 = double first cousins. (“Nibling” is a gender-neutral term for niece and nephew.)

Another way of visualizing how dispersal generates the observed pattern of isolation-by-distance is to plot the distribution of geographic distances separating pairs of individuals with different pedigree relationships (Fig 3A). Close relatives tend to be located closer geographically: for example, the distance between full-siblings is significantly less than the distance between pairs with *r* = 0.25 (half-siblings, grandparent-grandchildren, and aunt/uncle-niblings; Wilcoxon rank sum test, *p* = 0.01). More generally, if we compare a given pedigree relationship class (*r*) with the pedigree relationship class that is half as related (0.5r), we find shorter distances in the more related pairs for all sequential comparisons out to third cousins (comparing pairs with *r* to 0.5*r*, Wilcoxon rank sum test, *p* < 0.003 for all except for the comparison between *r* = 0.0625 and *r* = 0.03125; S4 Table). Geographic distances between two males with a close, known pedigree relationship are shorter than in either female-female or male-female comparisons (Fig 3A, S5 Table), and this pattern holds generally in comparisons up to second cousins.

**Fig 3.**
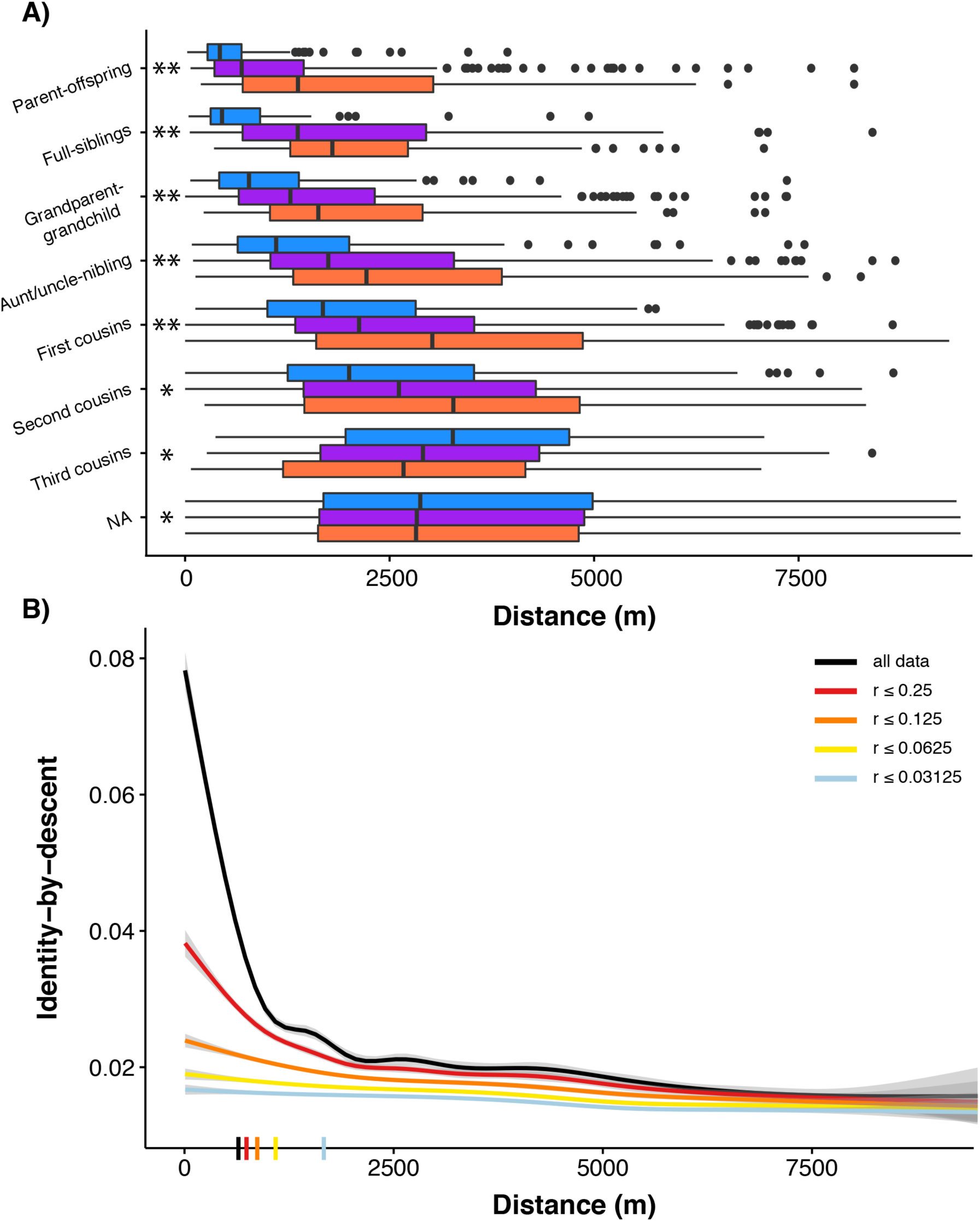
Distances between Florida Scrub-Jay individuals of close pedigree relatedness explains, in part, the observed isolation-by-distance patterns. (A) Distances between all possible male-male (blue), male-female (purple), and female-female (salmon) comparisons separated by pedigree relationship. Significant differences using the Kolmogorov-Smirnov Test are indicated with two asterisks when all three comparisons were significantly different (MM-FF, MM-MF, MF-FF) and a single asterisk when only MM-FF and MM-MF comparisons were significantly different. The distance between parent-offspring pairs is significantly shorter than the distance between full-siblings (Wilcoxon rank sum test, *p* = 5.20 x 10^−9^) and the distance between full-siblings is significantly shorter than the distance between pairs with *r* = 0.25 (Wilcoxon rank sum test, *p* = 0.01). (B) Loess curves of distance versus identity-by-descent in autosomal SNPs for all possible unique pairwise comparisons with separate lines showing sequential removal of pedigree relationship classes. The strength of isolation-by-distance decreases as highly related pairs are removed. δ values, the distance where identity-by-descent drops halfway to the mean (see text for details), are shown as dashes on the x-axis.

We can further assess the contribution of various relationship types by sequentially removing pedigree relationship classes and observing the resulting isolation-by-distance curves (Fig 3B). As expected, the relationship between identity-by-descent and geographic distance flattens and the strength of isolation-by-distance (measured by δ) decreases as closely related pairs are removed (Fig 3B, S6 Table). For example, removing pairs with *r* ≥ 0.5 (parent-offspring and full-siblings) and *r* ≥ 0.25 (parent-offspring, full-siblings, half-siblings, grandparents, and aunt/uncle-nibling) caused significant increases in δ (Fig 3B, S6 Table). However, even after removing all pairs with *r* ≥ 0.0625 we still see a significant pattern of isolation-by-distance (S6 Table). Therefore, isolation-by-distance is not driven only by highly related individuals. Instead, it appears that highly related individuals (*r* ≥ 0.25) play a primary role in determining the strength of the observed isolation-by-distance patterns (measured by δ), but isolation-by-distance still exists even when these individuals are removed from the dataset. The pattern of isolation-by-distance in more distantly related pairs suggests that isolation-by-distance is generated from dispersal events over many generations even at this small spatial scale, and is not simply a result of dispersal events over only one or two generations.

### Isolation-by-distance patterns are also present in Z-linked SNPs

Patterns of genetic diversity on the Z chromosome are expected to differ from those on the autosomes because of the difference in inheritance patterns and sex-specific dispersal behavior [40]. In birds, males are the homogametic sex (ZZ), while females are heterogametic (ZW). Thus, the Z chromosome spends two-thirds of its evolutionary history in males. In addition, the Z chromosome has a smaller effective population size compared with the autosomes [41]. These facts lead to two predictions: (1) Owing to the reduced effective population size of the Z chromosome, we expect to see higher identity-by-descent on the Z compared to the autosomes. (2) Because females disperse much farther than males in this system, we expect to find more isolation-by-distance in Z-linked SNPs than in autosomal SNPs [31,40].

We separately assessed patterns of isolation-by-distance in 277 Z-linked SNPs. PCA results for Z-linked markers are similar to those observed in autosomes. We found significant correlations for PC1 and PC2 with UTM northing, though correlations between PC2 and UTM northing were significant only for all breeders and male only comparisons (S3 Fig, S1 Table). To fairly compare autosomes and Z chromosomes, which differ in the number of SNPs present, we used unbiased estimates of identity-by-descent for Z-linked and autosomal SNP comparisons. These unbiased estimates do not undergo the final transformation step involved in the estimates of identity-by-descent used previously, and therefore are not bounded by 0 and 1 (see S1 Text for more details). Though these unbiased, unbounded estimates can take negative values, they make comparisons between autosome and Z datasets more straightforward. Bounding identity-by-descent estimates by 0 and 1 for the Z chromosome would generate upwardly biased estimates. Note that the autosomal identity-by-descent estimates are based on a larger set of SNPs and so values are similar between the bounded estimates and “unbiased” estimates. Therefore the bias is minimal and not a problem for the previous autosomal analyses.

Similar to autosomal SNPs, isolation-by-distance patterns in Z-linked SNPs are stronger in male-male comparisons (δ = 615 m, [592, 639]) than in either female-female (δ = 979 m, [673, 2048]) or male-female comparisons (δ = 637 m, [601, 674]; S4 Fig). In accordance with theoretical predications, mean identity-by-descent is higher for the Z chromosome (0.014, [0.013,0.015]) compared with the autosomes (0.0027, [0.0024,0.0030]; Fig 4). However, we do not find evidence for more isolation-by-distance on the Z chromosome: δ for Z-linked SNPs (647 m, [620, 677]) is not significantly different from δ for autosomal SNPs (621 m, [608, 633]; Fig 4). It is possible that we lack the power to estimate identity-by-descent on the Z chromosome accurately, given the small number of Z-linked SNPs available (277), which leads to more noise and uncertainty in the estimates of identity-by-descent on the Z chromosome and therefore a high variance in δ. This is consistent with the larger standard errors for the Z (Fig 4) and the larger confidence interval for δ on the Z. Future work will increase marker density on the Z to increase resolution and will incorporate maternally-inherited markers like the W and mitochondria to provide additional insights into the consequences of sex-biased dispersal on markers with different inheritance modes.

**Fig 4.**
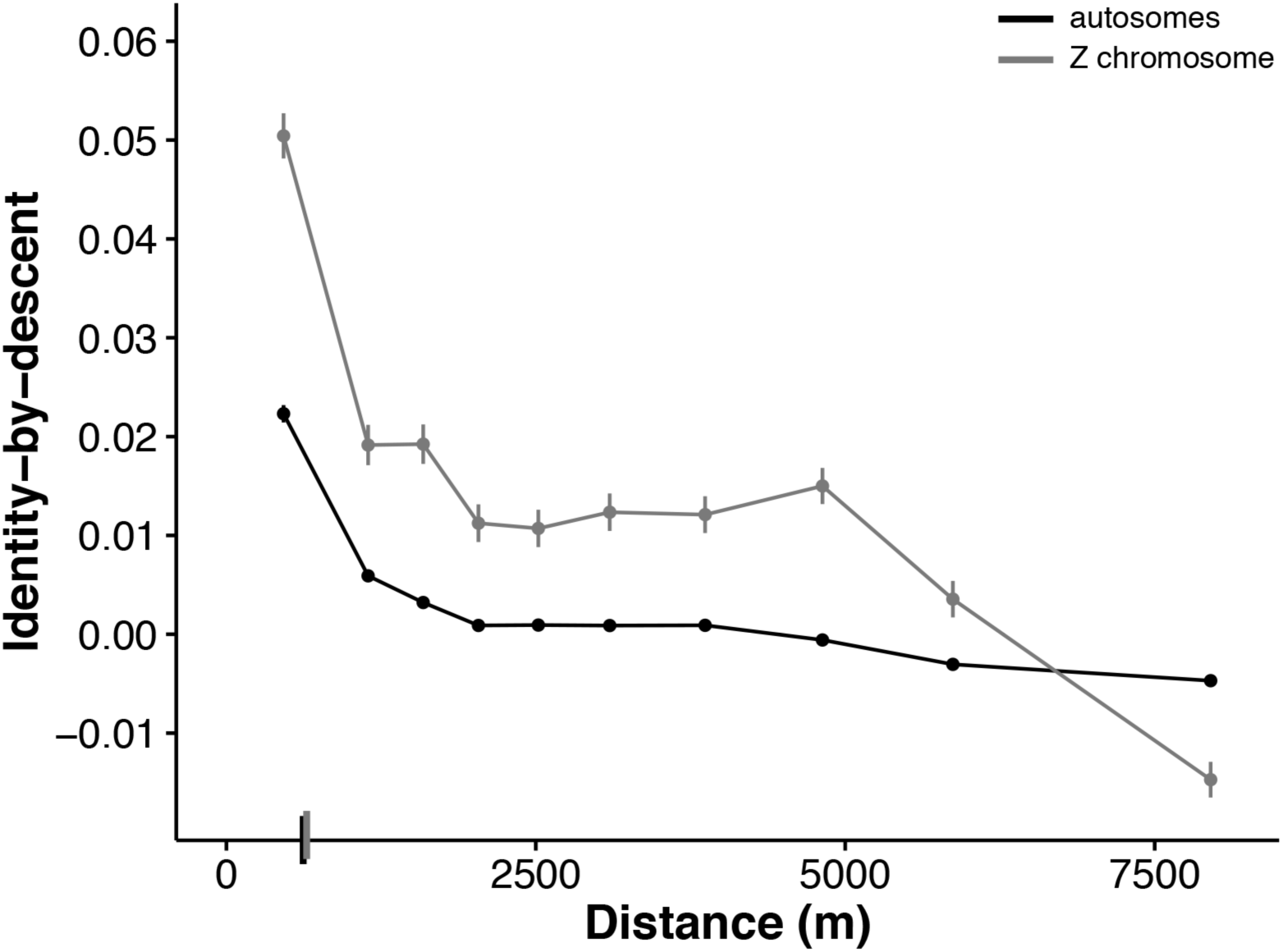
Isolation-by-distance in autosomal and Z-linked SNPs. Geographic distance versus unbiased identity-by-descent for autosomal (black) and Z-linked (gray) SNPs for all possible unique pairwise comparisons showing higher mean identity-by-descent in Z-linked SNPs (0.014) than in autosomal SNPs (0.0027). Here we use untransformed estimates of identity-by-descent to avoid biases introduced by the different numbers of autosomal and Z-linked SNPs (see text for details). Identity-by-descent values are binned across 10 distance quantiles and shown as mean ± SE. δ values, the distance where identity-by-descent drops halfway to the mean (see text for details), are shown as dashes on the x-axis.

### Simulations can reconstruct observed geographic structure

To test our understanding of the population mechanisms leading to fine-scale isolation-by-distance, we used simulations to explore whether observed patterns could be predicted strictly by dispersal curves and other population parameters. We first conducted simulations of local dispersal in a contiguous population to determine how well the observed distribution of geographic distances between individuals of known pedigree relationships was predicted by the observed natal dispersal curves. Assuming that the dispersal curves are constant and that dispersal distance has negligible heritability, we simulated the distance between individuals of a known, close pedigree relationship using random draws from the sex-specific dispersal curves. For example, for two female first cousins, we first simulated the dispersal distances of the parental siblings from the grandparental nest (randomly picking their sexes). We then simulated dispersal distances of the two female cousins from their respective parental nests and calculated the distance (*d*) between them (Fig 5A). We repeated this procedure 10,000 times to obtain a distribution of *d*.

**Fig 5.**
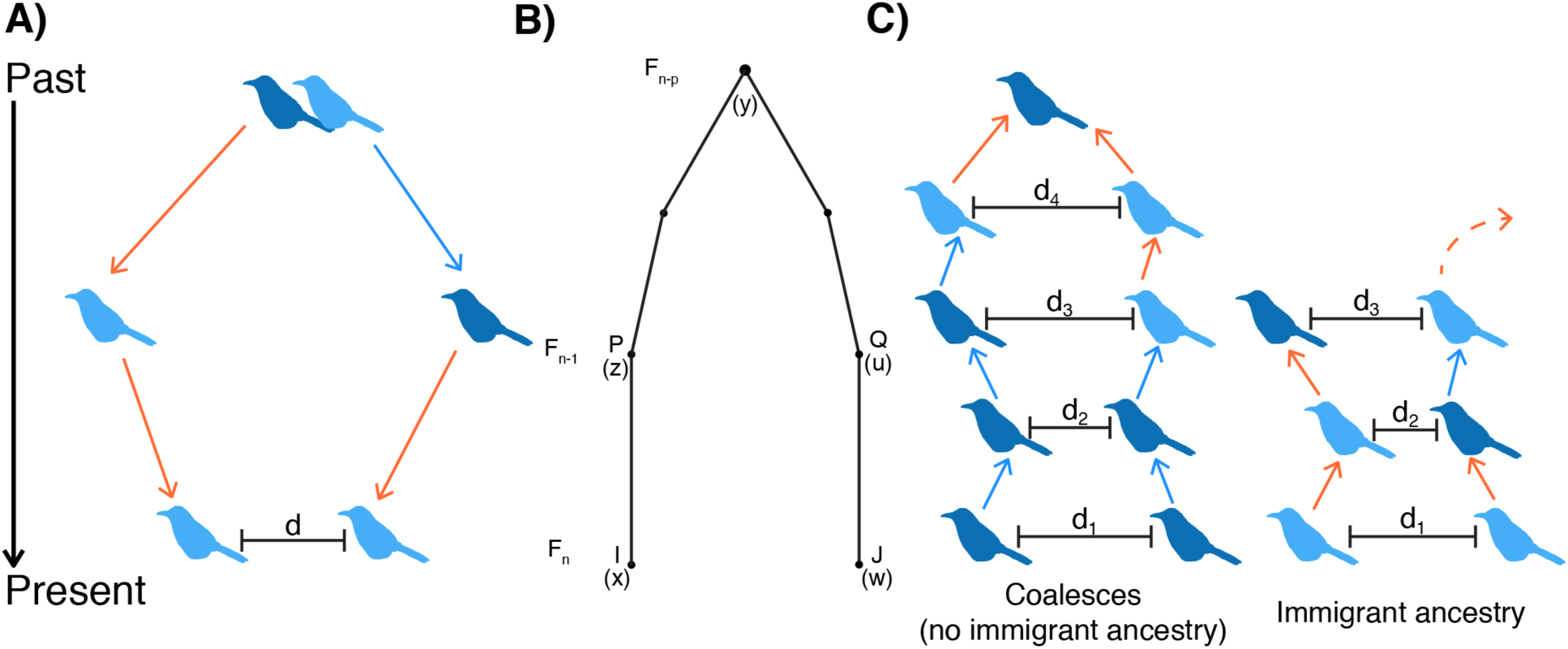
Overview of dispersal and coalescent simulations of isolation-by-distance. (A) An example schematic of a dispersal simulation for two female first cousins. Our simulations were over a two-dimensional space, but here we show dispersal on a onedimensional line for visualization purposes. For the dispersal simulations, we start with the most recent common ancestor for a pair of individuals of known pedigree relationship and simulate dispersal events forward in time until the present. In this case, we start at the grandparental nest, simulate dispersal distances (and angles) of the parents, and then dispersal of the two cousins. Light blue birds are females and dark blue are males. Arrows indicate male (blue) and female (salmon) dispersal events drawn from the dispersal curves. In most simulations, sexes of all ancestors are determined by a coin flip. (B) The gametic kinship chain from Malécot’s theory of isolation-by-distance. A locus from individual *I* born at location *x* and a locus from individual *J* born at location *w* in generation *F_n_* are identical-by-descent if both are descended from the same locus in their common ancestor in generation *F_n-p_* Under Malécot’s model, genetic relatedness of individuals should decrease as the distance between them increases. Redrawn from [9]. (C) Illustration of two possible outcomes in the coalescent simulations. In these simulations, we start with a pair of individuals of specified sex separated by distance *d_1_* and trace their ancestral lineages backwards in time until we either reach a common ancestor or one of the ancestors was an immigrant. In each generation, the probability a given pair coalesces is sampled directly from the pedigree. *M* is the probability a parental individual is an immigrant. Using empirical estimates of identity-by-descent between closely related pairs and immigrants, we generated expected identity-by-descent values for each pair.

We found that the dispersal simulations generally nicely reconstruct the observed distribution of geographic distances between related individuals up to second cousins (Fig 6, S7 Table; Kolmogorov-Smirnov Test with Bonferroni correction, *p* > 0.004 for most pairs). For more distantly related pairs, some of the simulations are significantly different from the observed distances (Fig 6, S7 Table; Kolmogorov-Smirnov Test, *p* < 0.004 for male-female first cousins and female-female second cousins). Notably, the observed distributions in male-male comparisons of closely related uncle-nephew pairs are significantly different from the simulated distributions – we see more short distances between individuals in the observed data than expected from the simulations (Fig 6, S7 Table).

**Fig 6.**
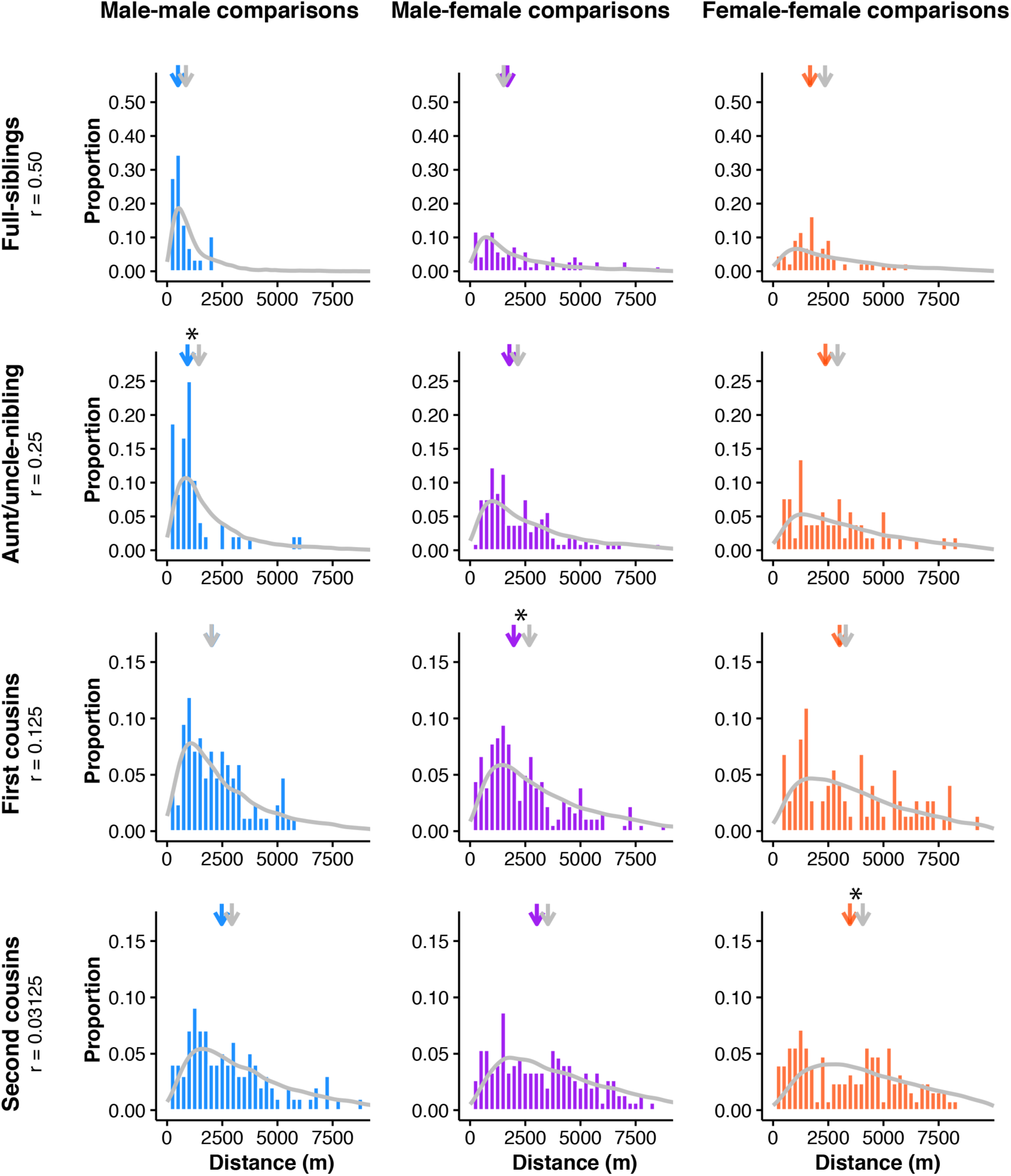
Dispersal simulations can reconstruct the observed distribution of geographic distances between related pairs. Simulated (gray line) and observed (colored histograms) dispersal values for full-sibling, aunt/uncle-nibling, first cousin, and second cousin comparisons. Male-male comparisons are shown in blue, male-female comparisons in purple, and female-female comparisons in salmon. Median values for the simulated (gray) and observed (colored) distributions are indicated by arrows above each plot. Simulated distributions that were significantly different from the observed distribution using the Kolmogorov-Smirnov Test are marked with asterisks above the median arrows.

The distance simulations described above randomized the sexes for all ancestral individuals and therefore averaged across all possible lineages for a given pedigree relationship. However, given the strongly sex-biased dispersal in the Florida Scrub-Jay, we expect the geographic distance between a given pair of individuals to also depend on the sexes of the ancestors. For example, two females can be cousins because their mothers are siblings (four female dispersal events), their mother and father are siblings (three female and one male dispersal events), or because their fathers are siblings (two female and two male dispersal events).

To assess the relationship between the sex of the ancestors and geographic distance between a pair of individuals of a given pedigree relationship, we conducted additional simulations of first cousins in which we fixed the sexes for the two common ancestors (aunts or uncles) in addition to the focal individuals (the cousins). As predicted, we found that the median geographic distance between two cousins strongly correlates with the number of female dispersal events in the lineage (Spearman rank correlation: *ρ* = 0.8208, *p* = 0.0067). For example, the median distance between two cousins depends on the number of female dispersal events in their lineage, such that male cousins related through their fathers (median ± SE = 1,715 ± 130 m) are geographically closer than male cousins related through their mothers (2,474 ± 235 m). Similar to our more general dispersal simulations (*i.e.,* those with randomized ancestral sexes), we found that the simulated distributions closely fitted the empirical patterns (S5 Fig, S8 Table). The observed distributions only differed from the simulated distributions in simulations with a male-female cousin pair related by their fathers (S8 Table; Kolmogorov-Smirnov Test, *p* = 0.0005).

In nature, we know that dispersal movements are largely restricted to the bounded area that is the study population. Because our natal dispersal curves include only within-population dispersal events, we do not think a violation of this assumption is problematic for simulations of closely related pairs, which involve just a few dispersal events. To accurately simulate distances between more distantly related pairs, we would need to consider the spatial extent of the population and not allow dispersal movements outside of population boundaries.

### Simulations can reconstruct observed genetic structure

Malécot envisioned identity-by-descent as being due to the chain of ancestry running from present day individuals back to their shared ancestors (“les chaînes de parenté gamétique”; Fig 5B; [9, 42]). These ideas are the forerunner of modern coalescent theory [43, 44]. Malécot’s interpretation of the relationship between identity-by-descent and geographic distance reflects the fact that geographically close pairs of individuals are more likely to be closely related, *i.e.,* trace back to a more recent common ancestor (coalesce), than geographically distant individuals [9].

To empirically demonstrate the underlying mechanisms behind Malécot’s model, we calculated the expected identity-by-descent values as a function of geographic distance for male-male, male-female, and female-female pairs using a spatially-explicit coalescent model. We parameterized these simulations using the observed pedigree, dispersal curves, immigration rate, and basic demographic information about the study system. We extended Malécot’s framework to include immigration from other populations because previous work has demonstrated a non-negligible rate of immigration into our study population [36]. For a given pair of individuals, we traced the ancestry of their two alleles at each autosomal locus backwards in time until the two lineages found a common ancestor or at least one of the lineages was a descendant of an immigrant into the population (Fig 5C). The probability that a lineage in a given generation was brought into the population by an immigrant (*M*) is given by the proportion of individuals who are immigrants. If one or both of our lineages traced back to an immigrant, we assigned the pair of individuals the observed level of identity-by-descent between immigrants. We kept track of the geographic location of each non-immigrant ancestor by sampling dispersal events from the natal dispersal curve. If our lineages are a distance *d_k_* apart in generation *k*, the probability of our lineages finding a shared ancestor in the next generation back (*k*+1) is given by the proportion of pairs that are *d_k_* apart who are full-siblings, half-siblings, or parent-offspring pairs (see S6 Fig). If the two lineages traced to one of these relationships, we assigned them the expected level of identity-by-descent for that relationship. We simulated expected identity-by-descent values for many pairs of individuals at a given distance bin.

We ran five different simulations to investigate how increasing the complexity of the model improved our fit to the observed isolation-by-distance patterns in male-male, male-female, and female-female pairs (S2 Text). We began with a model that used sex-averaged values for all parameters. This model (M0) explained a large proportion of the variance in mean identity-by-descent across geographic distance for male-female pairs (coefficient of determination *R*^2^ = 0.90) but not for male-male and female-female comparisons (*R*^2^ = 0.61 and −0.10, respectively; Fig 7, S7 Fig, Table 1). We then tried to improve the fit of our model by incorporating sex-specific parameters. First, we simulated dispersal back in time in a sex-specific manner by sampling from the male or female dispersal curve. Because of the strongly female-biased dispersal in Florida Scrub-Jays, the per-generation coalescent probability for females is greater at larger distance bins, and immigrants are more likely to be female [34, 36]. By allowing sex-specific dispersal (model M1), sex-specific coalescent parameters (model M2), and also sex-specific immigration parameters (model M3), our models more closely reconstructed the observed relationship between identity-by-descent and geographic distance (*R*^2^ = 0.88-0.90 for model M3; S7 Fig, Table 1). The fully sex-specific model overestimated identity-by-descent at longer distances for male-male pairs, which we hypothesized was a result of observed isolation-by-distance in the immigrants. By extending our model to account for variation in relatedness among immigrants with distance, our final pedigree-based simulations (model M4) recovered the observed pattern of isolation-by-distance for both autosomal and Z-linked loci, with slightly lower performance for female-female comparisons and for the Z chromosome (S8 Fig, Table 1). The fact that our simulations, which only span 10 generations, recovered the observed decrease in genomic relatedness within 10 km suggests that limited dispersal can generate isolation-by-distance over short timescales in this population.

**Figure 7.**
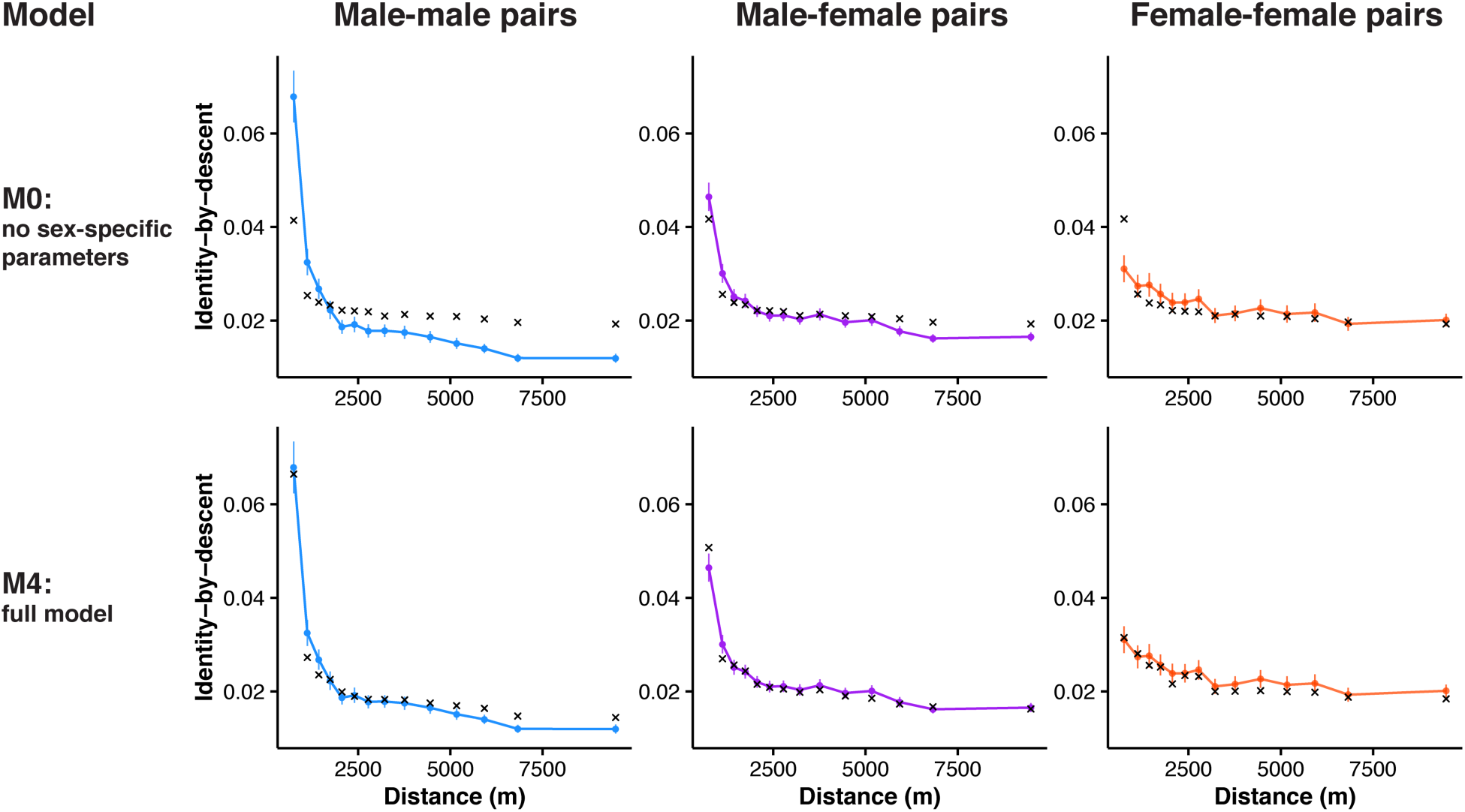
Coalescent simulations can reconstruct isolation-by-distance patterns. Simulated (black crosses) and observed (colored circles and line) autosomal isolation-by-distance patterns for male-male (blue), male-female (purple), and female-female comparisons (salmon). We ran five different simulations using the observed pedigree, dispersal curves, and immigration rate. Results are shown for two models: the simplest model with no sex-specific parameters (M0) on top and our final model with sex-specific parameters and isolation-by-distance in immigrants (M4) on bottom. By increasing the biological realism of our models, we can recover the observed pattern of isolation-by-distance. The coefficient of determination for the final model is 0.98 for male-male comparisons, 0.96 for male-female comparisons, and 0.78 for female-female comparisons. See Table 1 and S7 Fig for full results.

**Table 1.**
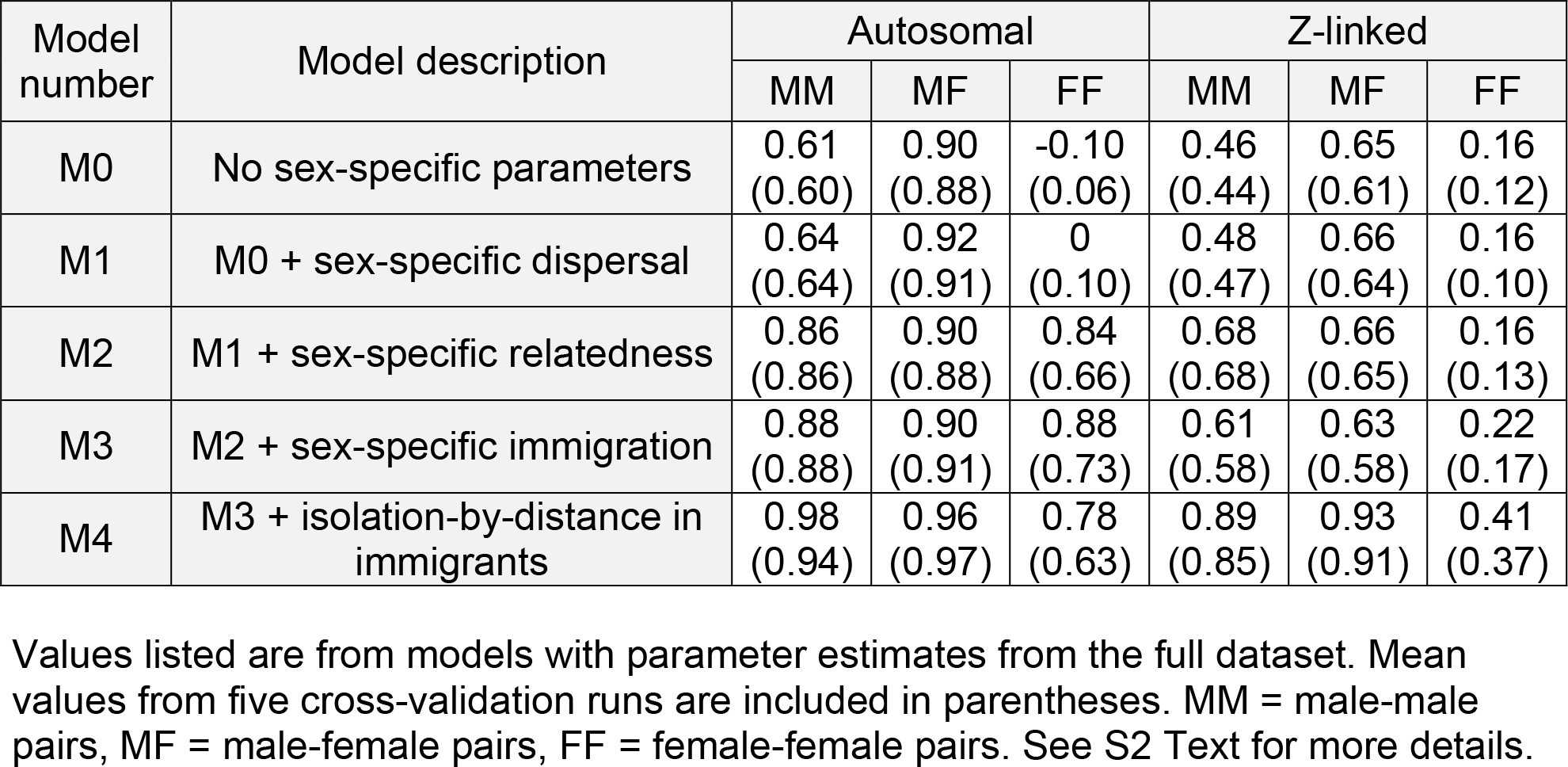
Coefficient of determination (*R^2^*) for different coalescent models for autosomal and Z-linked SNPs.

| Model number | Model description | Autosomal |  |  | Z-linked |  |  |
| --- | --- | --- | --- | --- | --- | --- | --- |
|  |  | MM | MF | FF | MM | MF | FF |
| M0 | No sex-specific parameters | 0.61<br>(0.60) | 0.90<br>(0.88) | -0.10<br>(0.06) | 0.46<br>(0.44) | 0.65<br>(0.61) | 0.16<br>(0.12) |
| M1 | M0 + sex-specific dispersal | 0.64<br>(0.64) | 0.92<br>(0.91) | 0<br>(0.10) | 0.48<br>(0.47) | 0.66<br>(0.64) | 0.16<br>(0.10) |
| M2 | M1 + sex-specific relatedness | 0.86<br>(0.86) | 0.90<br>(0.88) | 0.84<br>(0.66) | 0.68<br>(0.68) | 0.66<br>(0.65) | 0.16<br>(0.13) |
| M3 | M2 + sex-specific immigration | 0.88<br>(0.88) | 0.90<br>(0.91) | 0.88<br>(0.73) | 0.61<br>(0.58) | 0.63<br>(0.58) | 0.22<br>(0.17) |
| M4 | M3 + isolation-by-distance in immigrants | 0.98<br>(0.94) | 0.96<br>(0.97) | 0.78<br>(0.63) | 0.89<br>(0.85) | 0.93<br>(0.91) | 0.41<br>(0.37) |

Values listed are from models with parameter estimates from the full dataset. Mean values from five cross-validation runs are included in parentheses. MM = male-male pairs, MF = male-female pairs, FF = female-female pairs. See S2 Text for more details.

### Future directions

Here we have used single-marker estimates of genome-wide identity-by-descent to study relatedness. Additional power to infer recent demography and dispersal history can be gained by studying shared identity-by-descent blocks – linked segments of the genome that are shared identical-by-descent between pairs of individuals [45-47]. A number of methods exist for inferring identity-by-descent blocks from dense genotyping or sequencing data [48]. By tracing the spatial distribution of identity-by-descent blocks of varying lengths, we can uncover how recent dispersal shapes the transmission of genomic segments across the landscape. Furthermore, we will assess how dispersal shapes patterns of genetic variation over larger spatial scales by extending this approach to multiple populations spanning the entire range of this species. This question has vital conservation implications, as for example, decreasing rates of immigration are driving increased inbreeding depression within the population at Archbold Biological Station [36].

### Conclusion

Isolation-by-distance is a commonly observed pattern in nature. Despite its ubiquity and the frequent use of isolation-by-distance patterns to indirectly estimate dispersal in diverse organisms, few studies to date have deconstructed the causes of isolation-by-distance. Here, we have shown how limited dispersal can result in isolation-by-distance in the Florida Scrub-Jay. The extremely short dispersal distances of this species allow us to detect a signal of isolation-by-distance within a single, small contiguous population over just a few generations. In systems with longer dispersal distances, patterns of isolation-by-distance will likely only be observed over larger spatial scales, and reflect relatedness over potentially much longer timescales. The extensive dispersal, pedigree, and genomic data in this well-studied system provided a rare opportunity to empirically unpack and extend Malécot’s isolation-by-distance model [9]: we have shown how limited dispersal leads to closely related individuals being located closer together geographically, which results in a pattern of decreased genetic relatedness with increased geographic distance.

## Materials and Methods

### Study system: the Florida Scrub-Jay

The Florida Scrub-Jay is a cooperatively breeding bird endemic to Florida oak scrub habitat [32, 33]. Individuals live in groups consisting of a breeding pair and non-breeding helpers (often previous young of the breeding pair) within territories that are defended year-round. A population of Florida Scrub-Jays at Archbold Biological Station (Venus, Florida, USA) has been intensely monitored by two groups for decades: the northern half by Woolfenden, Fitzpatrick, Bowman, and colleagues since 1969 [32, 34] and the southern half by Mumme, Schoech, and colleagues since 1989 [35, 49]. Standard population monitoring protocols in both studies include individual banding of all adults and nestlings, mapping of territory size and location, and surveys to determine group composition, breeding status/success, and individual territory affiliation [32, 34]. Immigration into our study population is easily assessed because every individual is uniquely banded (so any unbanded individual is an immigrant). Blood samples for DNA have been routinely obtained from all adults and day 11 nestlings through brachial venipuncture since 1999. This intense monitoring has generated a pedigree of 14 generations over 46 years. All activities have been approved by the Cornell University and University of Memphis Institutional Animal Care and Use Committees and permitted by the US Geological Survey, the US Fish and Wildlife Service, and the Florida Fish and Wildlife Conservation Commission.

Here, we measured dispersal distances of individuals banded as nestlings within Archbold and that subsequently bred within Archbold between 1990 and 2013 (382 males and 290 females). We began our sampling in 1990 because the study site was expanded to its current size by 1990; hence, dispersal measures before this year are systematically shorter (*i.e.,* lack the longer distances). Thus, we have a comprehensive measure of dispersal tendencies of individuals within Archbold over a 24-year period. We measured natal dispersal distance as the distance from the center of the natal territory to the center of the first breeding territory in meters using ArcGIS Desktop v10.4 [50], independent of the age of first breeding (definition from [51]).

As part of a previous study, 3,984 individuals have been genotyped at 15,416 genome-wide SNPs using Illumina iSelect Beadchips [36]. Details of SNP discovery, genotyping, and quality control can be found in [36]. Here, we focused on breeding adults in Archbold during the years 2003, 2008, and 2013 (*n* = 513), when almost all individuals present have been genotyped. Autosomal SNPs were pruned for linkage disequilibrium using PLINK v1.07 [52]. We conducted analyses on both the entire set of SNPs and the dataset pruned for linkage disequilibrium. We found qualitatively similar results, so we present only the results from the pruned dataset here. Our final dataset included 7,843 autosomal and 277 non-pseudoautosomal Z-linked SNPs. All of the presented analyses were conducted on the combined dataset across all three years. For any individuals present in multiple years, we randomly selected presence in a single year for inclusion in this combined analysis.

### Relatedness measures

To determine genetic relatedness, we estimated the proportion of the genome shared identical-by-descent relative to the population frequency for all individual pairwise comparisons within and across years using the ‘genome’ option in PLINK v1.07 [52] for autosomal SNPs. As PLINK does not calculate identity-by-descent for sex-linked markers, we used a custom R script to estimate the proportion of the genome shared identical-by-descent for Z-linked SNPs (S1 File). Identity-by-descent for Z-linked SNPs was calculated using a method-of-moments approach using observed allele counts similar to that in [52]. Identity-by-descent values reported by PLINK are constrained to biologically plausible values between 0 and 1 in a final transformation step. To avoid introducing biases when comparing identity-by-descent estimates obtained from very different numbers of SNPs (on the Z chromosome versus the autosomes), we used untransformed autosomal and Z-linked identity-by-descent values for comparisons between the autosomes and Z. All identity-by-descent calculations used allele frequencies from the sample of all individuals in the population through time. See S1 Text for further details and S1 File for the R code.

Additionally, we estimated relatedness of all individual pairwise comparisons using the pedigree. We calculated the coefficient of relationship by using the ‘kinship’ function within the package kinship2 [53] in R v3.2.2 [54] and multiplied the values by two (to convert them from kinship coefficients). The pedigree-based coefficient of relationship was calculated separately for expectations under autosomal and Z-linked scenarios using the ‘chrtype’ option within the ‘kinship’ function. Because kinship2 assumes an XY system, we swapped the sex labels of our individuals and swapped mothers and fathers in the pedigree to calculate the coefficient of relationship for a ZW system. The autosomal coefficient of relationship *r* and proportion of the genome shared identical-by-descent are highly correlated (S9 Fig; Pearson’s product moment correlation: *t* = 688.85, *p* < 0.0001). Because genomic estimators of relatedness are more precise than pedigree-based estimators [55], we only report results for genomic measures of relatedness in the text (but see S10 Fig, S11 Fig, and S2 Table for analyses using pedigree-based measures of relatedness).

### Isolation-by-distance in genetic and pedigree data

We used three approaches to test for isolation-by-distance patterns in our data. First, we conducted principal component analysis on the autosomal and Z-linked genomic data using custom Perl and R scripts. We conducted separate analyses on males only, females only, and all individuals. We then compared the first two PC axes from each analysis with the UTM northing values of the territory centroids for each individual using Spearman rank correlations. To ensure these patterns were not driven by differences in genetic diversity within the study site, we estimated observed heterozygosity and inbreeding coefficients *(F^III^* from [56]) from the autosomal SNPs in PLINK. We compared individual heterozygosity and inbreeding coefficients with UTM northing and found no relationship (Pearson’s product-moment correlation, t = 1.493, *p* = 0.136 for heterozygosity, t = −1.559, *p* = 0.120 for inbreeding coefficient).

Second, we conducted Mantel correlogram tests using the ‘mantel.correlog’ function in the vegan package [57] in R v3.2.2 [54]. Mantel tests compare two distance matrices and test for significance through permutation of the matrix elements [58, 59]. While Mantel tests are useful for assessing linear relationships, they will not accurately represent the spatial structure found in systems with exponential-like decreases in structure (*i.e.,* strong spatial structure in the short distance classes that decreases and stabilizes at larger distances). Mantel correlograms are able to assess these more complex spatial structures by utilizing the traditional Mantel test within distinct distance bins [38, 39]. Here, we use Mantel correlograms to compare a matrix of individual pairwise comparisons of geographic distances to a matrix of pairwise comparisons of relatedness between individuals (either estimated from the genomic data or from the pedigree, and for autosomes or the Z chromosome). We conducted separate analyses for comparisons between males only, females only, and all individuals. Note that we cannot conduct Mantel correlograms on male-female comparisons alone, as we cannot use unbalanced matrices in this type of analysis. We limited our analyses to the following distance class bins to ensure that enough comparisons fell within each bin: 250-750 m, 750-1250 m, 1250-1750 m, 1750-2250 m, 2250-2750 m, 2750-3250 m, 3250-3750 m, 3750-4250 m, 4250-4750 m, 4750-5250 m. We did not include comparisons between breeders in the same territory or self-self comparisons (distance < 250 m). We performed 10,000 permutations to obtain corrected *p*-values.

Finally, we fitted a loess curve to the scatterplot of identity-by-descent and geographic distance between pairs of individuals. We tested for isolation-by-distance by determining whether identity-by-descent at the smallest distance interval was larger than the overall mean. To measure the strength of isolation-by-distance, we estimated the distance where identity-by-descent drops halfway to the mean from its maximum value, which we define as δ. To assess uncertainty in these estimates, we used a bootstrapping method in which we randomly resampled pairs with replacement, fitted a loess curve, and estimated identity-by-descent at distance bin 0, mean identity-by-descent, and δ. We repeated this procedure 1,000 times to obtain 95% bias-corrected and accelerated bootstrap confidence intervals.

### Dispersal simulations

We used simulations to determine whether we could generate the observed distribution of geographic distances between related pairs using only the natal dispersal curve. For each of several focal pairwise relationships (full-siblings, aunt/uncle-nibling, first cousins, and second cousins), we simulated dispersal events starting at their common ancestral nest and then recorded the resulting distance between the two focal individuals using a custom script in R (Fig 5A, S2 File). We located the shared ancestral nest of the birds at (0,0) in an unbounded two-dimensional habitat. The number of dispersal events for a given focal pair ranged from two (full-siblings) to six (second cousins). For each dispersal event, we randomly sampled a dispersal angle (0-360°) and a dispersal distance from the sex-specific dispersal distribution (Fig 1A). The sexes of the final individuals in the focal pair were fixed (either male-male, male-female, or female-female). In most cases, the sexes of ancestral individuals up to the common ancestor were chosen randomly with a coin flip. To further assess the impact of sex-specific dispersal on the distribution of geographic distances between pairs, we performed simulations for first cousins with fixed sexes for the two focal individuals (the cousins) and the two common ancestors (aunts or uncles). This resulted in nine possible simulations (with sex combinations of male-male, male-female, and female-female for both the focal pair and the common ancestors). We performed these simulations 10,000 times for each focal pairwise relationship, calculating the resulting distance between the two focal individuals each time. We determined the empirical distances between individuals of different pedigree relationships and compared the observed distributions to the simulated distributions using Kolmogorov-Smirnov tests and the means using Wilcoxon rank sum tests, both with Bonferroni corrections. Code for the dispersal simulations is included in S2 File.

### Coalescent simulations

We generated the expected isolation-by-distance pattern for the autosomes and the Z chromosome given the observed dispersal curves and immigration rate using spatially-explicit pedigree-based simulations that are extensions of Malécot’s model of isolation-by-distance [9]. For each pair of individuals, we simulated their lineages backwards in time until we reached a common ancestor or one or more of the lineages was a descendent of an immigrant into the population (Fig 5C). In each generation g, we first sampled a dispersal distance from the empirical sex-specific dispersal curve (Fig 1A) and a dispersal angle (0-360°) uniformly at random and calculated the geographic distance *d_g_* between the two individuals. Here, we assumed that there is no genetic variation for dispersal distance, and sampled a dispersal distance for all individuals of a given sex from the same distribution. After the first dispersal event, we randomly assigned sexes for all ancestors. We then calculated the probability that the two lineages located at distances (*d_1_*,…, *d_g_*) did not coalesce (share a common ancestor) or have an immigrant ancestor in the previous *g – 1* generations and the probability that they either coalesce or have an immigrant ancestor in generation g. Given the relatively small population size and high immigration rate, we found that nearly all pairs either shared a common ancestor or had an immigrant ancestor within 10 generations, and so we used *g* ≤ 10 (increasing this limit had no effect on our results). Here we define the probability that two individuals share a common ancestor in the preceding generation as the probability the pair is closely related (parent-offspring, full-siblings, or half-siblings). For a pair of individuals at distance d, we estimated the probability they are parent-offspring (*P_p_*(*d*)), full-siblings (*P_f_*(*d*)), or half-siblings (*P_h_*(*d*)) from the observed pedigree and distances between these relative classes (S6 Fig). We calculated the sex-specific probability an individual is an immigrant as the proportion of breeding male or female individuals in a given year who were not born in Archbold (*M* = 0. 197 for males and 0.345 for females). Using mean identity-by-descent values for immigrant-immigrant and immigrant-resident pairs obtained from our data, we estimated the expected proportion of the genome shared identical-by-descent for a given pair of individuals as follows:

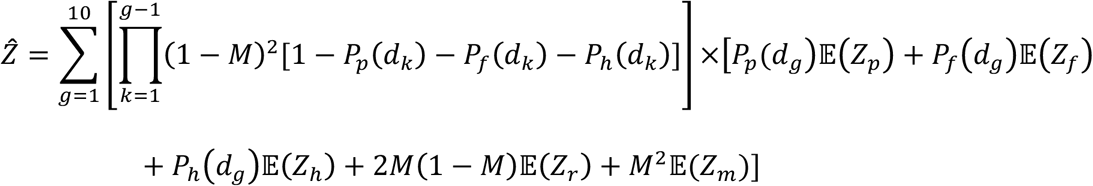

Where E(*Z_p_*), E(*Z_f_*), and E(*Z_h_*) are the expected identity-by-descent values for parent-offspring, full-sibling, and half-sibling pairs, respectively (S9 Table). E(*Z_m_*), and E(*Z_r_*) are the sex-specific empirical mean identity-by-descent values for immigrant-immigrant and immigrant-resident pairs, respectively. Because we found a pattern of isolation-by-distance in immigrant-immigrant pairs, we used expected identity-by-descent values for immigrant-immigrant and immigrant-resident pairs conditional on distance. We binned distances into 15 quantiles and ran 1,000 simulations for each distance bin. To evaluate the fit of our model, we calculated the coefficient of determination *R^2^* for each type of comparison as follows:

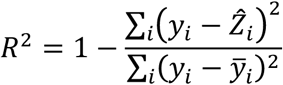

Where *y_i_* is the mean observed identity-by-descent value in distance bin *i* and *Z_i_* is the mean simulated identity-by-descent value in distance bin *i*. Note that it is possible to obtain negative values of *R^2^* when the model performs so poorly that the mean of the data provides a better fit than our model. We ran simulations using parameters estimated from the full dataset, and then performed two-fold cross-validation to check for over-fitting. As results from both sets of models were similar, we discuss results from the full dataset in the text. See S2 Text for the full derivation of our model and S3 File for the R code.

## Acknowledgments

We thank the many students, interns, and staff at Archbold Biological Station and the University of Memphis who collected the dispersal data. We thank the members of the Harrison, Clark, Lovette, and Coop labs for thoughtful comments on this manuscript. In memory of Richard G. Harrison without whom this collaboration would not have been formed.

## Supporting Information

**S1 Dataset. Dispersal data used in this study**

**S2 Dataset. Genetic and pedigree relatedness data used in this study**

**S1 File. R code for estimating identity-by-descent for Z-linked markers**

**S2 File. R code for dispersal simulations**

**S3 File. R code for coalescent simulations**

**S1 Text. Identity-by-descent estimation for Z-linked markers**

**S2 Text. Derivation of coalescent model**

## References

1. Clobert J, Danchin E, Dhont AA, Nichols JD. Dispersal. Oxford, UK: Oxford University Press; 2001.

2. Coulon A, Fitzpatrick JW, Bowman R, Lovette IJ. Mind the gap: genetic distance increases with habitat gap size in Florida scrub jays. Biol Lett. 2012;8(4):582–5. doi: 10. 1098/rsbl.2011. 1244. PMID: 22357936.

3. Coulon A, Fitzpatrick JW, Bowman R, Lovette IJ. Effects of habitat fragmentation on effective dispersal of Florida scrub-jays. Conserv Biol. 2010;24(4):1080–8. doi: 10.1111/j.1523-1739.2009.01438.x. PMID: 20151985.

4. Duckworth RA, Badyaev AV. Coupling of dispersal and aggression facilitates the rapid range expansion of a passerine bird. Proc Natl Acad Sci USA. 2007;104(38):15017–22. doi: 10.1073/pnas.0706174104. PMID: 17827278.

5. Wright S. Isolation by distance under diverse systems of mating. Genetics. 1946;31:39–59.

6. Wright S. Isolation by Distance. Genetics. 1943;28:114–38.

7. Malécot G. Les Mathématiques de l'Hérédité. Paris: Masson; 1948.

8. Malécot G. Heterozygosity and relationship in regularly subdivided populations. Theor Popul Biol. 1975;8:212–41.

9. Malécot G. Génétique des populations diploïdes naturelles dans le cas d'un seul locus. III. Parenté, mutations et migration. Ann Génét Sél Anim. 1973;5(3):333–61.

10. Vekemans X, Hardy OJ. New insights from fine-scale spatial genetic structure analyses in plant populations. Mol Ecol. 2004;13(4):921–35. doi: 10.1046/j.1365- 294X.2004.02076.x.

11. Meirmans PG. The trouble with isolation by distance. Mol Ecol. 2012;21(12):2839–46. doi: 10.1111/j.1365-294X.2012.05578.x.

12. Koenig WD, Van Vuren D, Hooge PN. Detectability, philopatry, and the distribution of dispersal distances in vertebrates. Trends Ecol Evol. 1996;11(12):514–7.

13. Broquet T, Petit EJ. Molecular Estimation of Dispersal for Ecology and Population Genetics. Annu Rev Ecol Evol Syst. 2009;40(1):193–216. doi: 10.1146/annurev.ecolsys.110308.120324.

14. Epperson BK. Estimating dispersal from short distance spatial autocorrelation. Heredity. 2005;95(1):7–15. doi: 10.1038/sj.hdy.6800680. PMID: 15931252.

15. Rousset F. Genetic differentiation and estimation of gene flow from F-statistics under isolation by distance. Genetics. 1997;145:1219–28.

16. Li MH, Merila J. Genetic evidence for male-biased dispersal in the Siberian jay (*Perisoreus infaustus*) based on autosomal and Z-chromosomal markers. Mol Ecol. 2010;19:23–5281. doi: 10.1111/j.1365-294X.2010.04870.x. PMID: 20977509.

17. Slatkin M. Gene flow in natural populations. Annu Rev Ecol Syst. 1985;16:393–430.

18. Excoffier L, Foll M, Petit RJ. Genetic Consequences of Range Expansions. Annu Rev Ecol Evol Syst. 2009;40(1):481–501. doi: 10.1146/annurev.ecolsys.39.110707.173414.

19. Leighton GM, Echeverri S. Population genomics of Sociable Weavers *Philetairus socius* reveals considerable admixture among colonies. J Ornithol. 2016;157:483–92. doi: 10.1007/s10336-015-1307-1.

20. Garroway CJ, Radersma R, Sepil I, Santure AW, De Cauwer I, Slate J, et al. Fine-scale genetic structure in a wild bird population: the role of limited dispersal and environmentally based selection as causal factors. Evolution. 2013;67(12):3488–500. doi: 10.1111/evo.12121. PMID: 24299402.

21. Temple HJ, Hoffman JI, Amos W. Dispersal, philopatry and intergroup relatedness: fine-scale genetic structure in the white-breasted thrasher, *Ramphocinclus brachyurus*. Mol Ecol. 2006;15(11):3449–58. doi: 10.1111/j.1365-294X.2006.03006.x. PMID: 16968282.

22. Foerster K, Valcu M, Johnsen A, Kempenaers B. A spatial genetic structure and effects of relatedness on mate choice in a wild bird population. Mol Ecol. 2006;15(14):4555–67. doi: 10.1111/j.1365-294X.2006.03091.x. PMID: 17107482.

23. Quaglietta L, Fonseca VC, Hájková P, Mira A, Boitani L. Fine-scale population genetic structure and short-range sex-biased dispersal in a solitary carnivore, *Lutra lutra*. J Mammal. 2013;94(3):561–71. doi: 10.1644/12-mamm-a-171.1.

24. Weber JN, Bradburd GS, Stuart YE, Stutz WE, Bolnick DI. Partitioning the effects of isolation by distance, environment, and physical barriers on genomic divergence between parapatric threespine stickleback. Evolution. 2016. doi: 10.1111/evo. 13110. PMID: 27804120.

25. van Dijk RE, Covas R, Doutrelant C, Spottiswoode CN, Hatchwell BJ. Fine-scale genetic structure reflects sex-specific dispersal strategies in a population of sociable weavers (*Philetairus socius*). Mol Ecol. 2015;24(16):4296–311. doi: 10.1111/mec. 13308. PMID: 26172866.

26. Dobson FS. Competition for mates and predominant juvenile male dispersal in mammals. Anim Behav. 1982;30:1183–92.

27. Greenwood PJ. Mating systems, philopatry and dispersal in birds and mammals. Anim Behav. 1980;28:1140–62.

28. Goudet J, Perrin N, Waser P. Tests for sex-biased dispersal using bi-parentally inherited genetic markers. Mol Ecol. 2002;11:1103–14.

29. Roy J, Gray M, Stoinski T, Robbins MM, Vigilant L. Fine-scale genetic structure analyses suggest further male than female dispersal in mountain gorillas. BMC Ecol. 2014;14:21.

30. Prugnolle F, de Meeus T. Inferring sex-biased dispersal from population genetic tools: a review. Heredity. 2002;88:161–5. doi: 10.1038/sj/hdy/6800060.

31. Segurel L, Martinez-Cruz B, Quintana-Murci L, Balaresque P, Georges M, Hegay T, et al. Sex-specific genetic structure and social organization in Central Asia: insights from a multi-locus study. PLoS Genet. 2008;4(9):e1000200. doi: 10.1371/journal.pgen.1000200. PMID: 18818760.

32. Woolfenden GE, Fitzpatrick JW. The Florida Scrub Jay: Demography of a Cooperative-breeding Bird. Princeton, New Jersey: Princeton University Press; 1984.

33. Woolfenden GE, Fitzpatrick JW. Florida Scrub-Jay (Aphelocoma coerulescens). Poole A, editor. Ithaca: Cornell Lab of Ornithology; 1996.

34. Fitzpatrick JW, Woolfenden GE, Bowman R, editors. Dispersal distance and its demographic consequences in the Florida scrub-jay. Proceedings of the XXII International Ornithological Congress; 1999; Durban: South Africa.

35. Schoech SJ, Mumme RL, Moore MC. Reproductive endocrinology and mechanisms of breeding inhibition in cooperatively breeding Florida scrub jays (*Aphelocoma c. coerulescens*). Condor. 1991;93:354–64.

36. Chen N, Cosgrove Elissa J, Bowman R, Fitzpatrick John W, Clark Andrew G. Genomic consequences of population decline in the endangered Florida scrub-jay. Curr Biol. 2016. doi: 10.1016/j.cub.2016.08.062.

37. Paradis E, Baillie SR, Sutherland WJ, Gregory RD. Patterns of natal and breeding dispersal in birds. J Anim Ecol. 1998;67:518–36.

38. Borcard D, Legendre P. Is the Mantel correlogram powerful enough to be useful in ecological analysis? A simulation study. Ecology. 2012;93(6):1473–81.

39. Diniz-Filho JAF, Soares TN, Lima JS, Dobrovolski R, Landeiro VL, de Campos Telles MP, et al. Mantel test in population genetics. Genet Mol Biol. 2013;36(4):475–85.

40. Schaffner SF. The X chromosome in population genetics. Nat Rev Genet. 2004;5(1):43–51. doi: 10.1038/nrg1247. PMID: 14708015.

41. Charlesworth B. Effective population size and patterns of molecular evolution and variation. Nat Rev Genet. 2009;10(3):195–205. doi: 10.1038/nrg2526. PMID: 19204717.

42. Ishida Y. Sewall Wright and Gustave Malécot on isolation by distance. Philos Sci. 2009;76(5):784–96.

43. Slatkin M, Veuille M. Modern developments in theoretical population genetics: the legacy of Gustave Malécot: Oxford University Press; 2002.

44. Nagylaki T. Gustave Malécot and the transition from classical to modern population genetics. Genetics. 1989;122(2):253–68.

45. Ralph P, Coop G. The geography of recent genetic ancestry across Europe. PLoS Biol. 2013;11(5):e1001555. doi: 10.1371/journal.pbio.1001555. PMID: 23667324.

46. Ringbauer H, Coop G, Barton NH. Inferring recent demography from isolation by distance of long shared sequence blocks. Genetics. 2017.

47. Baharian S, Barakatt M, Gignoux CR, Shringarpure S, Errington J, Blot WJ, et al. The Great Migration and African-American Genomic Diversity. PLoS Genet. 2016;12(5):e1006059. doi: 10.1371/journal.pgen.1006059.

48. Browning SR, Browning BL. Identity by descent between distant relatives: detection and applications. Annu Rev Genet. 2012;46:617–33. doi: 10.1146/annurev-genet-110711-155534. PMID: 22994355.

49. Mumme RL. Do helpers increase reproductive success? Behav Ecol Sociobiol. 1992;31:319–28.

50. ESRI. ArcGIS Desktop: Release 10. Redlands, CA: Environmental Systems Research Institute; 2011.

51. Greenwood PJ, Harvey PH. The natal and breeding dispersal of birds. Annu Rev Ecol Syst. 1982;13:1–21. doi: 10.1146/annurev.es.13.110182.000245.

52. Purcell S, Neale B, Todd-Brown K, Thomas L, Ferreira M, Bender D, et al. PLINK: a toolset for whole-genome association and population-based linkage analysis. Am J Hum Genet. 2007;81:559–75.

53. Therneau TM, Sinnwell J. kinship2: Pedigree Functions. 2015.

54. R Core Team. R: A language and environment for statistical computing. Vienna, Austria: R Foundation for Statistical Computing; 2015.

55. Kardos M, Luikart G, Allendorf FW. Measuring individual inbreeding in the age of genomics: marker-based measures are better than pedigrees. Heredity. 2015;115(1):63–72. doi: 10.1038/hdy.2015.17. PMID: 26059970.

56. Yang J, Lee SH, Goddard ME, Visscher PM. GCTA: a tool for genome-wide complex trait analysis. Am J Hum Genet. 2011;88(1):76–82. doi: 10.1016/j.ajhg.2010.11.011. PMID: 21167468.

57. Oksanen J, Blanchet FG, Kindt R, Legendre P, Minchin PR, O'Hara RB, et al. vegan: Community Ecology Package. 2015.

58. Mantel N. The detection of disease clustering and a generalized regression approach. Cancer Res. 1967;27:209–20.

59. Mantel N, Valand RS. A technique of nonparametric multivariate analysis. Biometrics. 1970;26(3):547–58.

60. Grossman M, Eisen EJ. Inbreeding, coancestry, and covariance between relatives for X-chromosome loci. J Hered. 1989;80:137–42.

